# New Lipophilic Fluorescent Dyes for Exosome Labeling: Monitoring of Cellular Uptake of Exosomes

**DOI:** 10.1101/2020.02.02.931295

**Authors:** Takashi Shimomura, Ryo Seino, Kaori Umezaki, Asako Shimoda, Takatoshi Ezoe, Munetaka Ishiyama, Kazunari Akiyoshi

**Author notes:** **Corresponding Author** Takashi Shimomura.

## Abstract

We have developed three types of exosomal membrane binding fluorescent probes, Mem Dye-Green, Mem Dye-Red and Mem Dye-Deep Red, to monitor exosome uptake into cells. The dyes contain a cyanine group as a fluorescent scaffold, which allows for highly sensitive fluorescence imaging of the exosome. These dyes can also be used to observe the dynamics of exosomes in live cells. The use of PKH dyes (Figure 1), which are currently the most widely-used fluorescent probes for exosome labeling, has some limitations. For example, PKH dyes tend to aggregate to form exosome-like nanoparticles, and these nanoparticles are uptaken by cells. Moreover, Mehdi suggested that the use of PKH dyes triggers an enlargement of the exosome size owing to membrane fusion or intercalation. To overcome the limitations of PKH dyes, we introduce amphiphilic moieties to the cyanine. To investigate the characteristics of the Mem Dyes as exosome labeling probes, we perform nanoparticle tracking analysis (NTA), zeta potential measurement and confocal microscopy. The Mem Dyes show excellent performance for exosome labeling (no aggregation and less size shift).

## Introduction

Exosomes are small membrane vesicles (30–150 nm in diameter) secreted by all cell types and can be found in almost all biofluids and in cell culture media^1^. Exosomes mediate intercellular communication by exchanging proteins, DNA, RNA and lipids between donor and recipient cells, and modulate the biological activities in target cells through receptor-ligand interactions^2^. Exosomes secreted by cells are uptaken through different routes and mechanisms, which include fusion with the plasma membrane and a range of endocytic pathways such as receptor-mediated endocytosis, phagocytosis, lipid raft-dependent endocytosis, micropinocytosis, and macropinocytosis^3^. Many researchers have studied exosome uptake into target cells and the *in vivo* biodistribution by tracking the exosomes. Among a range of exosome tracking approaches, fluorescent lipid membrane (lipophilic) dyes, such as PKH26^4^, PKH67^5^, DiI^6^, and DiD^7^, have been widely used. However, as Mehdi reported, dye-to-dye aggregation likely occurs during the labeling process when PKH dyes are used^8,9^. It was further shown that aggregated PKH was uptaken by the cells and such PKH cannot be distinguished from PKH-labeled exosomes. This phenomenon of PKH may cause false positives in microscopy imaging. It is considered that this phenomenon of PKH arises from a less established structural design (e.g., poor hydrophilicity and weak anchoring ability).

Aware of these issues of PKH, we have employed poly-ethyleneglycol to improve the hydrophilicity and introduce a negatively charged moiety to a symmetric two-armed cyanine. Further, to allow the newly developed fluorescent probe to be used in as many applications as possible, we attempt to synthesize three probes that emit different colors (green, red, deep red) and name them as Mem Dyes. When the Mem Dye was used for fluorescent labeling of purified exosomes, no dye-to-dye aggregation was observed in aqueous buffer. Moreover, the Mem Dyes themselves did not significantly change the size and population of exosomes whereas PKH did. As we have suggested above, it is very important that the exosome labeling fluorescent probes should not change any exosome characteristics (e.g., size and population). Therefore, the newly developed Mem Dyes are promising alternatives for use in exosome tracking.

## Results

### Nanoparticle Tracking Analysis (NTA)

Mem Dyes and PKH dyes in aqueous solution were analyzed by NTA (LM10-HSBFT 14, Nanosight) to investigate the generation of aggregates. Dyes were dissolved in DMSO and diluted to 10 μmol/L in phosphate-buffered saline. No aggregation was observed in the experiments with Mem Dyes, although PKH dyes produced dye-to-dye aggregates (100–500 nm size) (Figure 2).

**Figure 1.**
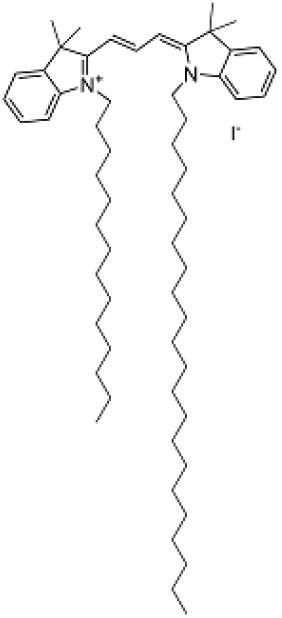
Chemical structure of PKH26.

**Figure 2.**
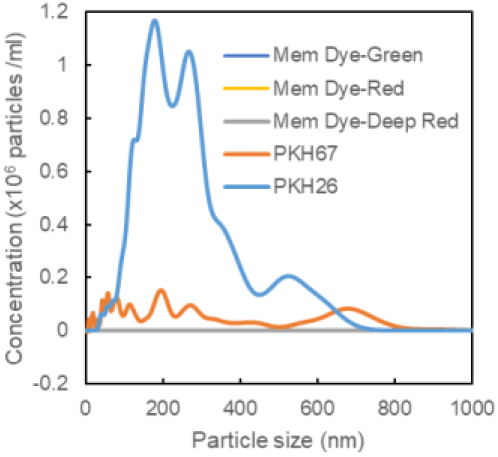
Nanoparticle tracking analysis of the amount and size distribution of dye aggregates.

Next, we stained exosomes with the Mem Dyes and PKH dyes to determine the effect of the dyes on the physical properties of the exosomes. As shown in Figure 3, Mem Dyes did not change the particle size and population of the exosomes. Conversely, the PKH dyes stained exosome sample contained a much lower nanoparticle population in a heterogeneous size distribution (Figure 3).

**Figure 3.**
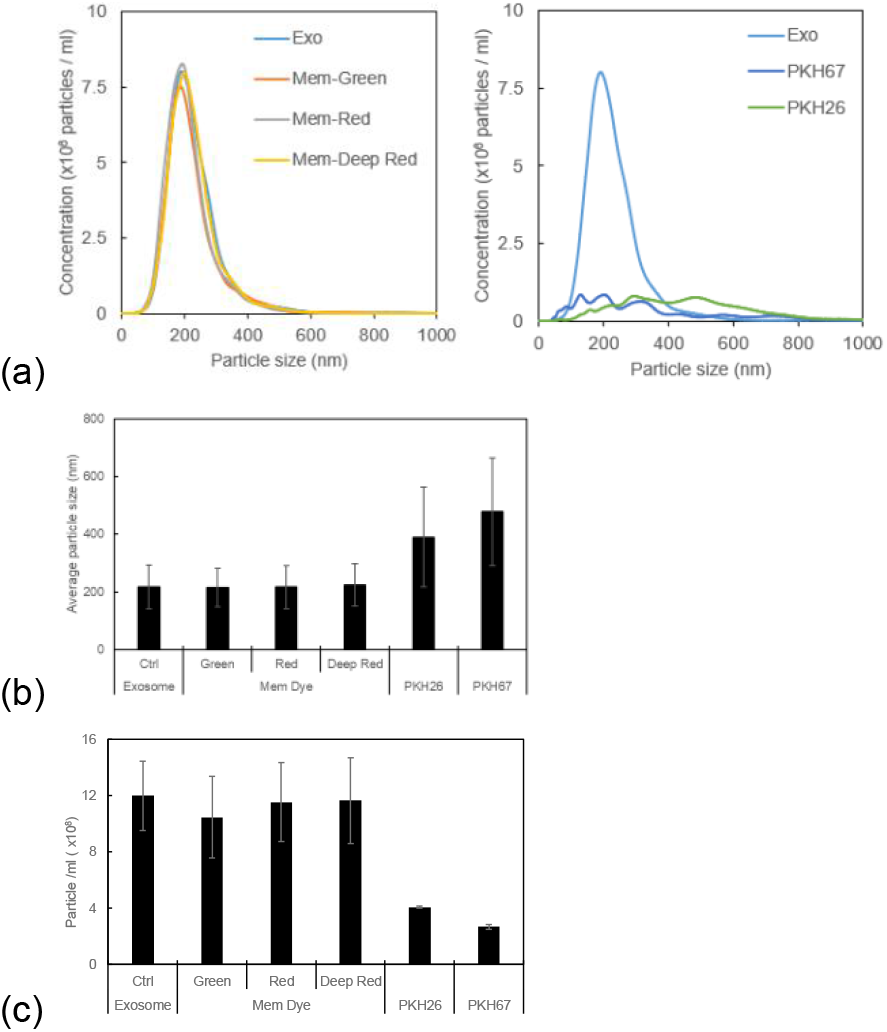
(a) Nanoparticle tracking analysis of the size distribution of dye-stained or unstained exosomes. (b) The average size of stained and unstained exosomes. (c) Particle counts of dye-stained or unstained exosomes analyzed by NTA.

### Zeta potential determination

Zeta potentials of Mem Dye and PKH dye stained exosomes were measured using a Zetasizer Nano ZSP (Malvern, PA, USA). Briefly, exosomes were stained with each dye at a dye concentration of 10 μmol/L. Then, the excess dye was removed by ultrafiltration and the residues were resuspended with 100 mmol/L HEPES solution. The zeta potential of the stained exosome samples was measured. As a negative control, the zeta potential of an exosome was also analyzed. As shown in Figure 4, PKH dye-stained exo-somes have lower zeta potential than Mem Dye-stained and unstained exosomes. The zeta potential difference between PKH dye-stained and Mem Dye-stained exosome might result from the structural difference (PKH dyes: positively charged, Mem Dyes: negatively charged).

**Figure 4.**
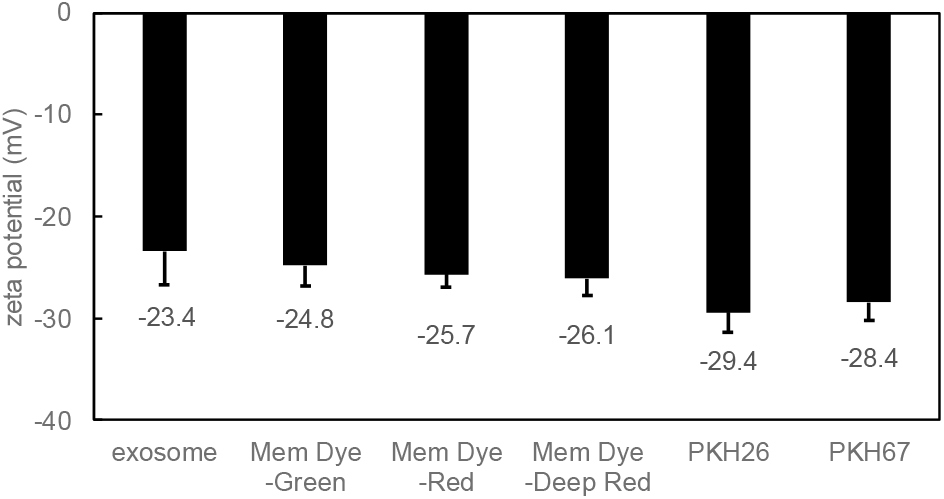
Zeta potential analysis of exosomes.

### Monitoring of exosome uptake in HeLa cells

Mem Dyes or PKH dyes stained exosomes were added to HeLa cells and the uptake/distribution of the stained exo-somes were monitored under a fluorescence microscope. All dyes clearly exhibited fluorescent puncta and no fluorescent signal was observed from the outside of cells. Interestingly, whereas larger puncta were detected in the PKH dye stained samples, Mem Dyes showed smaller puncta (Figure 5). It is considered that the larger puncta we observed in the staining with PKH dye were dye-to-dye aggregates as Mehdi suggested.

**Figure 5.**
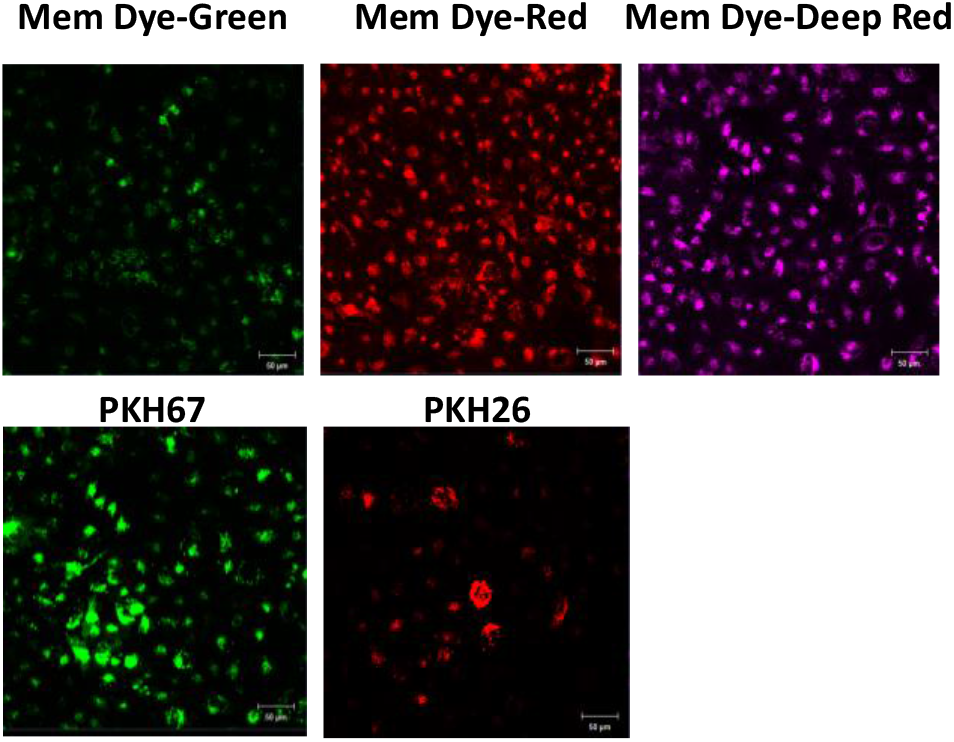
Imaging of labeled exosomes in HeLa cells.

## Discussion

PKH67 and PKH26 are widely used for exosome monitoring. They have lipophilic carbocyanine as a fluorescent scaffold and aliphatic tails that allow them to intercalate with the lipid bilayers of the exosome. As previous findings demonstrated, we also observed the limitations of PKH dyes by NTA and microscopy analysis. According to our NTA experiment, the larger nanoparticles found in the experiment with PKH may result from either nanoparticle linking or enlargement of the nanoparticles by staining. We assume the different results between Mem Dyes and PKH dyes were dependent on the chemical structure of each dye. As described in the introduction, the Mem Dyes had a hydrophilic and negatively charged moiety introduced. The Mem Dyes did not form dye-to-dye aggregates and did not change the population and size of the exosomes during labeling. Moreover, when we compared the exosome uptake into HeLa cells, fluorescent spots were observed from both the Mem Dyes and PKH dyes. However, the larger fluorescent spots, which were attributed to dye aggregates, were not observed when using Mem Dyes. From these results, we conclude that this structural modification from PKH dyes allows for the exact monitoring of exosomes achieved by our experiments. Here we have reported some zeta potential measurement results. However, further experiments should be performed because the exosome uptake efficiency depends on the zeta potential of nanoparticles^11^.

### Summary

Mem Dyes, which are newly developed fluorescent probes, are non-aggregatable and do not change the particle size of an exosome during experiments. Exosomes stained with Mem dyes in mammalian cells can be identified more clearly compared with those stained with PKH dyes without changing the zeta potential. Exosomes are generating considerable interest as drug carriers, potent medicine candidates and diagnostic markers. However, exosomes are not well-understood so far owing to their characteristics, heterogeneous size and internal components (proteins, nucleic acids, membrane and so on). We believe our findings reported here will allow exosome research to be further advanced.

## Notes

The authors declare no competing financial interests.

